# The kinase occupancy of T-cell coreceptors reconsidered

**DOI:** 10.1101/2022.08.01.502332

**Authors:** Alexander M. Mørch, Falk Schneider, Edward Jenkins, Ana Mafalda Santos, Scott E. Fraser, Simon J. Davis, Michael L. Dustin

**Affiliations:** Kennedy Institute of Rheumatology, Roosevelt Drive, University of Oxford, Oxford, OX3 7FY, United Kingdom; MRC Weatherall Institute of Molecular Medicine, John Radcliffe Hospital, University of Oxford, Oxford, OX3 9DS, United Kingdom; Translational Imaging Center, University of Southern California, Los Angeles, California 90089, United States of America

**Keywords:** Coreceptor, Lck, stoichiometry, T-cell signaling, FCS, diffusion

## Abstract

The sensitivity of the αβ T-cell receptor (TCR) is enhanced by the coreceptors CD4 and CD8αβ, which are expressed primarily by cells of the helper and cytotoxic T-cell lineages, respectively. The coreceptors bind to major histocompatibility complex (MHC) molecules and associate intracellularly with the Src-family kinase Lck, which catalyzes TCR phosphorylation during receptor triggering. Although coreceptor-kinase occupancy was initially believed to be high, a recent study suggested that most coreceptors exist in an Lck-free state, and that this low occupancy helps to effect TCR antigen discrimination. Here, using the same method, we found instead that the CD4-Lck interaction was stoichiometric (~100%) and that the CD8αβ-Lck interaction was also substantial (~60%). We confirmed our findings in live cells using fluorescence cross-correlation spectroscopy (FCCS) to measure coreceptor-Lck co-diffusion *in situ*. After introducing structurally guided mutations into the intracellular domain of CD4, we used FCCS to show that stoichiometric Lck coupling required an amphipathic α-helix present in CD4 but not CD8α. In double-positive cells expressing equal numbers of both coreceptors, but limiting amounts of kinase, CD4 out-competed CD8αβ for Lck. In T cells, TCR signaling induced CD4-Lck oligomerization but did not affect the high levels of CD4-Lck occupancy. These findings help settle the question of kinase occupancy and suggest that the binding advantages that CD4 has over CD8 could be important when Lck levels are limiting.

**Significance statement:** CD4 and CD8αβ are archetypal coreceptor proteins that potently enhance T-cell antigen sensitivity but how they function is still debated. A fundamental question that remains incompletely resolved is: what fractions of the coreceptors bind the signal-initiating kinase, Lck? Using *in vitro* assays and non-invasive fluorescence fluctuation spectroscopy in live cells, we show that most coreceptors are occupied by Lck at the surface of live cells. The structural basis for important differences in the kinase occupancy of CD4 and CD8αβ is also identified. These results provide important context for refining current models of both TCR antigen recognition and cell fate decisions made during thymopoiesis.

## Introduction

Conventional αβ T cells are divided into two major functional subsets depending on which of two coreceptors, CD4 or CD8, they express, that recognize class II and class I major histocompatibility complex (MHC) proteins, respectively. CD4^+^ T cells provide ‘help’ to antibody-producing B cells by secreting cytokines whereas CD8^+^ T cells are directly cytotoxic^1^. The coreceptors increase the sensitivity of T cell receptor (TCR) signaling through recruitment of the Src-family kinase Lck, which is especially important for the recognition of low affinity antigens^2,3^. One explanation for this effect is that TCR triggering is a two-step process requiring initial incipient phosphorylation of the TCR by free Lck, which is very quickly followed by the recruitment of a coreceptor-Lck complex. This second step is thought to occur through bidentate interactions of the coreceptor-Lck complex with the TCR (via Lck) and its pMHC ligand, which further enhances receptor phosphorylation^4,5^. A second possibility is that, rather than increasing the sensitivity of signaling, the coreceptors effect antigen discrimination. According to this point of view, which was prompted by measurements by Stepanek *et al*. suggesting that kinase occupancy is very low^6^, binding of cognate pMHC to the TCR is followed by processive coreceptor ‘scanning’ until Lck is co-recruited, favoring the phosphorylation of long-lived TCR complexes.

An important differentiator between these proposals is the extent to which coreceptors are bound to Lck, but the level of kinase occupancy is not yet agreed^7^. Early co-immunoprecipitation (co-IP) experiments showed that CD4-Lck occupancy is high (~90%) in primary T cells^8^, in line with evidence from bioluminescence resonance energy transfer (BRET) experiments which indicate that CD4 is invariably bound to Lck at the surface of HEK293 cells^9^ (**Figure 1a**). The recent proposal by Stepanek and colleagues that CD4 is only fractionally coupled to Lck (~6%) in thymocytes according to flow cytometric co-IP (FC-IP) assays was, therefore, unexpected^6^ (**Figure 1b**).

**Figure 1.**
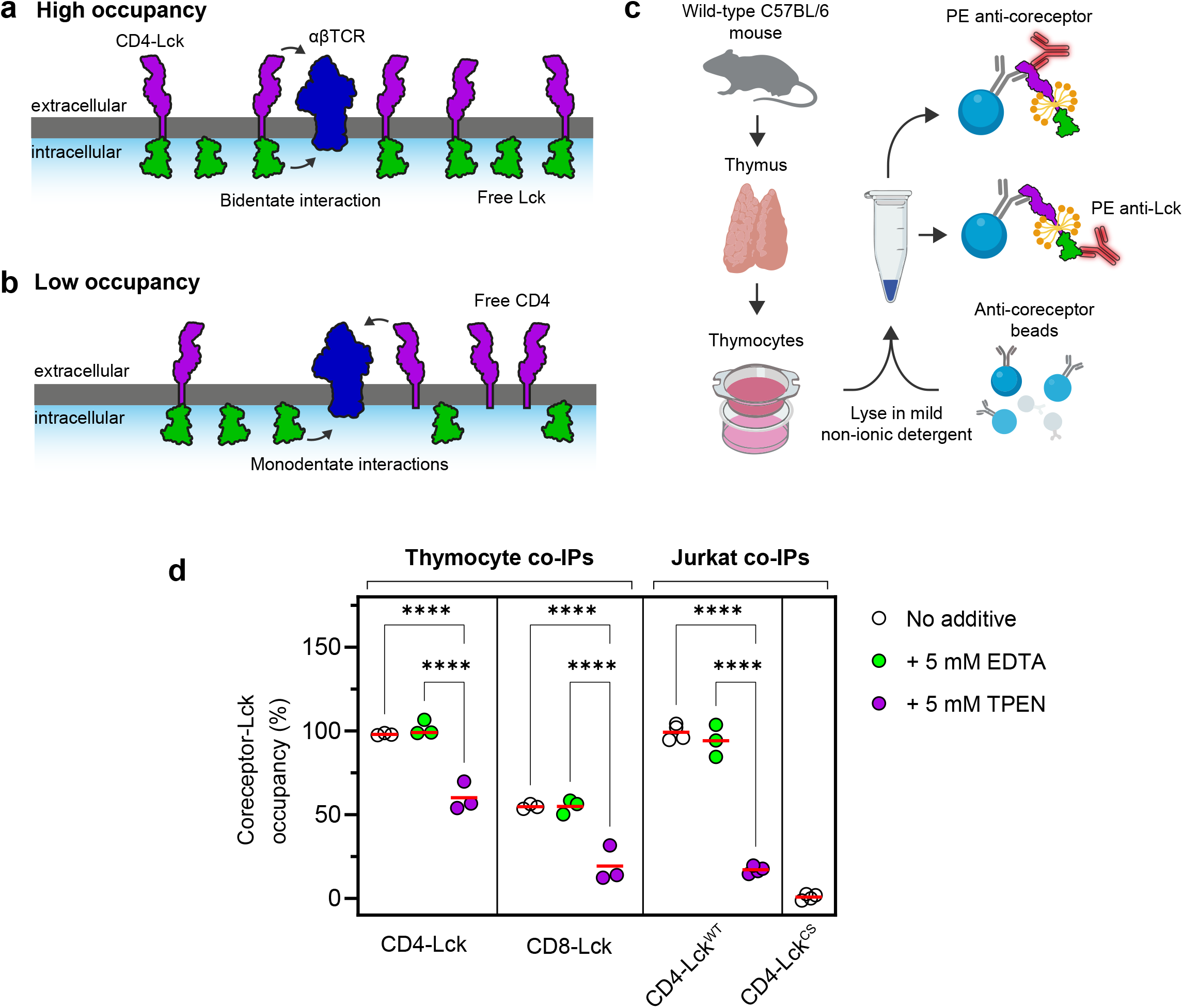
High coreceptor-Lck occupancy in both thymocytes and Jurkat T cells is only partially disrupted by chelation. **a**. and **b**. Depiction of high or low coreceptor occupancy at the T cell surface. At high occupancy (**a**), CD4-Lck complexes can form bidentate interactions with TCR-pMHC via Lck binding to phosphorylated ITAMs and CD4 binding to MHC-II (not depicted). At low occupancy (**b**), CD4 and Lck bind independently to TCR-pMHC complexes. Protein models based on crystal structures of CD4, Src and αβTCR (PDB IDs: 1WIQ, 3EL8 and 6JXR, respectively). **c**. Schematic of the FC-IP assay used in this study, created with BioRender. **d**. FC-IP measurements of coreceptor-Lck occupancy in bulk thymocytes and Jurkats expressing murine (m)CD4 and mLck (see **Figure S1c**) indicate that CD4 and CD8αβ form high-occupancy complexes with Lck independently of EDTA. TPEN partially dissolves the interaction but there is a residual TPEN-resistant component in both thymocytes and Jurkat T cells. Each circle indicates a flow cytometry measurement from either one mouse or a replicate measurement of the indicated Jurkat cell line. 10^6^ cells were lysed for each experiment. Clasp-deficient Lck contains the C20S and C23S mutations (Lck^CS^). Significance testing was performed with a one-way ANOVA followed by Tukey’s correction for multiple comparisons, and only significant comparisons are shown. \*\*\*\**P* < 0.0001.

Here we report that, in the course of reproducing the recent FC-IP experiments, CD4 and CD8αβ were observed to have a high rather than low kinase occupancy in both primary thymocytes and transfected Jurkat T cells. To confirm these findings *in situ*, we used scanning fluorescence correlation spectroscopy (sFCCS)^10,11^ to measure the co-diffusion of coreceptors and surface-bound Lck in live cells. We found that CD4 and Lck associate at a high occupancy (~100%) at the cell surface, and that this efficient coupling depends both on the conserved Zn^2+^ clasp motif and the amphipathic helix in the intracellular domain (ICD) of CD4^12^. CD8αβ lacks an amphipathic helix and, accordingly, exhibits a lower Lck occupancy (~60%). Consistent with this difference, when the coreceptors were co-expressed in HEK293T cells with limiting amounts of kinase, CD4 outcompetes CD8αβ for Lck. Lastly, we present evidence that CD4-Lck occupancy is unaffected by early TCR signaling, indicating that coreceptor-Lck complexes are stable and persist beyond receptor triggering.

## Results

### FC-IPs indicate high coreceptor-Lck occupancy in thymocytes and Jurkat T cells

CD4-Lck occupancy measurements (**Figure 1a, 1b**) have traditionally been performed using co-IP assays in buffers that contain chelating agents, such as ethylenediaminetetraacetic acid (EDTA), to inhibit metalloprotease activity^6,8,13^. However, EDTA has also been shown to disrupt the Zn^2+^-dependent association of CD4 and Lck^14^ which might explain some of the discrepancies between reports. To determine whether the presence of Zn^2+^ chelators could affect the measured occupancy, we performed the FC-IP assay developed and used by Stepanek *et al*. (**Figure 1c, Figure S1a**)^6^. Thymocyte lysates from wild-type C57Bl/6 mice were incubated with beads conjugated to anti-mouse (m)CD4 antibodies before staining for either mCD4, mouse (m)Lck or rat CD48 as a negative control (**Figure 1d**). Residual bead fluorescence is measured with flow cytometry, and the background-subtracted fluorescence ratio between Lck and CD4 offers a measure of coreceptor-Lck coupling ratio. Since the majority of thymocytes are CD4^+^ CD8αβ^+^ double-positive (DP)^15^, we expected this assay to produce a low CD4-Lck occupancy (6%-17%) as reported in experiments with sorted DP thymocytes^6,16^. Instead, however, we found that the interaction was effectively saturated (*i*.*e*., all CD4 molecules were bound to Lck) (**Figure 1d, Figure S1b**). The addition of EDTA to the lysates had no effect, implying that the detergent-solubilized complexes are resistant to Zn^2+^ chelation under these conditions. Pre-treatment of cells with millimolar amounts of a membrane-permeable Zn^2+^ chelator, *N,N,N*′,*N*′-tetrakis(2-pyridinylmethyl)-1,2-ethanediamine (TPEN), which was also expected to disrupt the CD4-Lck interaction *in vitro*^17,18^ reduced the occupancy by ~40% (**Figure 1d**). CD8αβ was substantially occupied by Lck (~55%) and this was reduced in the presence of TPEN treatment of live cells, but not EDTA treatment of lysates. This confirmed that sequestering Zn^2+^ *in situ* could disrupt the coreceptor-Lck interaction, but the residual association of precipitated proteins suggested either incomplete Zn^2+^ chelation by TPEN or the Zn^2+^-independent association of coreceptors and Lck captured in detergent micelles during lysis.

To test if the partial dissociations were due to non-specific interactions, we used lentiviral gene delivery to transduce mCD4 and mLck into Jurkat T cells in which expression of endogenous CD4/Lck had been abolished using CRISPR-Cas9 (**Figure S1c**). We observed a high CD4-Lck occupancy for the wild-type interaction irrespective of EDTA treatment but TPEN substantially dissociated the complex (**Figure 1d**). When the Zn^2+^ clasp of Lck was mutated (Lck^C20S,C23S^ = Lck^CS^), no Lck precipitated, consistent with previous data showing that the clasp is essential for binding^19^. These results lead us to conclude that TPEN was only partially chelating Zn^2+^ in the FC-IP assay, but indicated that the high occupancies observed in thymocytes were not an artefact of the *in vitro* solubilization method. However, the recent reports of low kinase occupancy^6^ cannot be explained by the presence of EDTA in the solubilization buffer. To confirm our observations, we sought non-invasive approaches to measure the stoichiometry of coreceptor-Lck complexes at the surface of live cells.

### Analysis of CD4 and Lck occupancy in live cells

To measure CD4-Lck occupancy at the surface of individual cells we employed fluorescence cross-correlation spectroscopy (FCCS), a two-color fluctuation spectroscopy method developed to quantify molecular dynamics *in situ*^20,21^ (**Figure 2a**). Extraction and fitting of autocorrelation functions (ACFs) provides information on diffusion and density, while cross-correlation functions (CCFs) can be used to measure heterotypic interactions (*i*.*e*., between spectrally distinct species)^11^. In order to perform FCCS of membrane proteins, which diffuse slowly and are prone to photobleaching, we acquired fluorescence signals in scanning-mode (sFCCS) which scans multiple contiguous pixels sequentially to improve signal-to-noise and reduce photobleaching effects (**Figure S2a**)^22^. To this end, full-length human CD4 and Lck constructs were linked to FCCS-appropriate tags (CD4-mCherry2 and Lck-mEGFP, ref. ^23^) and expressed in HEK293T cells grown on glass coverslips (**Figure 2b**). Intensity fluctuations were acquired using a confocal microscope in photon-counting mode and correlation analyses were performed using FoCuS_scan software^24^ (**Figure 2c**). Using the calibrated size of the observation volume, we also determined fluorophore density (see **SI Materials and Methods**) and confirmed that all constructs were expressed at physiological levels similar to the normal densities of CD4 and Lck (**Figure S2b**). To control for underestimation of interactions that might result from non-fluorescent FPs, we generated a positive control consisting of the transmembrane domain of CD4 linked covalently to both mEGFP and mCherry2 (‘Tandem’, **Figure 2c**). Co-diffusion of mEGFP and mCherry2, *i*.*e*., the cross-correlation quotient (*q*, see **SI Materials and Methods**), was then determined as the ratio between the CCF amplitude and the minimum ACF amplitude^21^.

**Figure 2.**
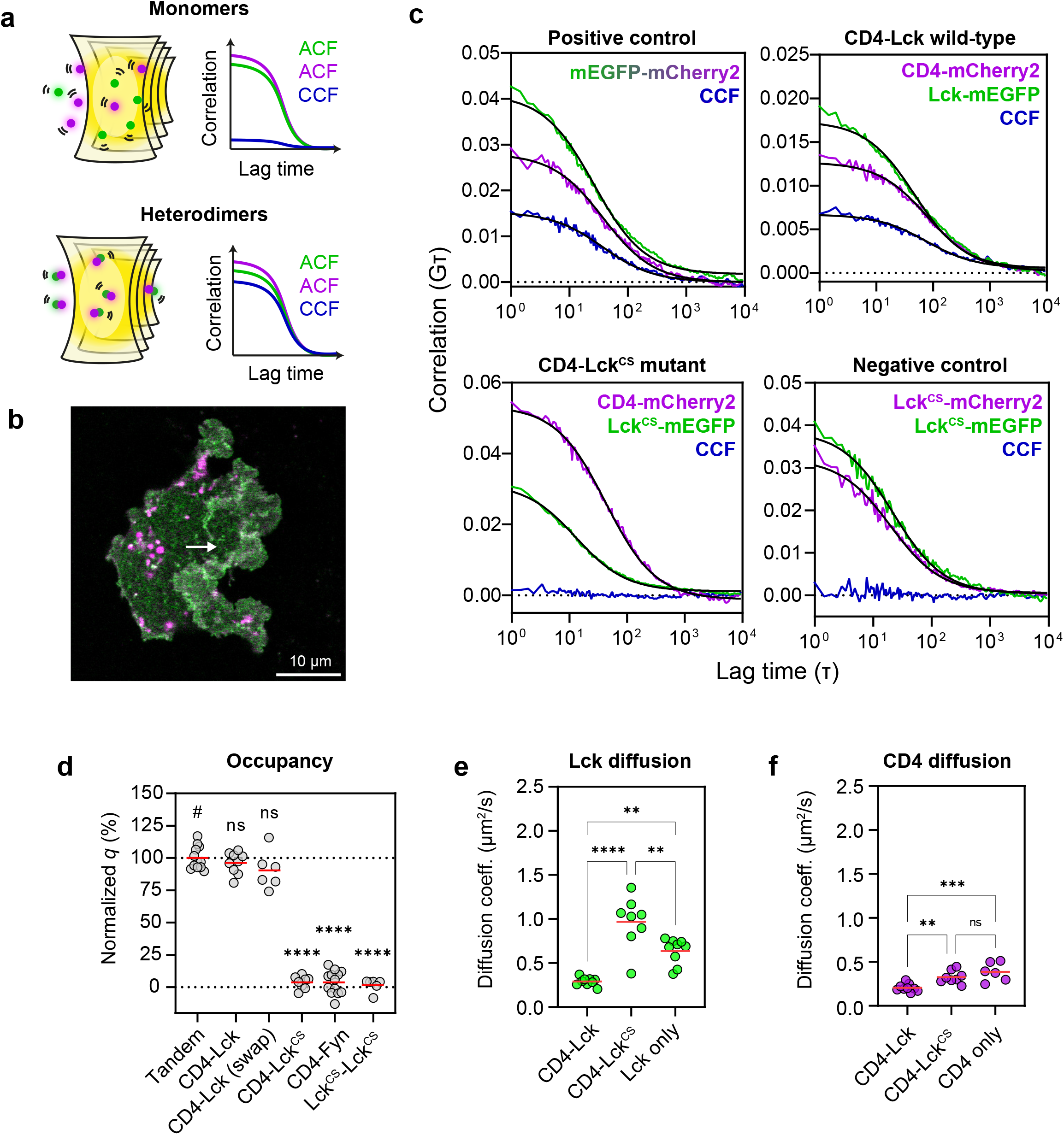
Scanning-FCCS measurements show a high CD4-Lck occupancy. **a**. Schematic of scanning-FCCS (sFCCS) volumes and correlation curves from monomeric (top) and dimeric (bottom) species. The autocorrelation functions (ACFs, magenta and green) are identical for monomers and dimers and can be fitted to extract the diffusion terms. The cross-correlation curves (CCFs, blue) are different between monomers and dimers with a low amplitude (top) indicating no co-diffusion and a high amplitude (bottom) indicating complete co-diffusion. **b**. Representative confocal image of a HEK293T cell co-transfected with CD4-mCherry2 and Lck-mEGFP acquired in photon-counting mode. sFCCS measurements are acquired in a similar fashion with shorter pixel dwell times to sample fluorescence fluctuations. The length and position of a typical sFCCS line-scan is depicted by the white arrow. **c**. Representative line-averaged ACFs (magenta/green) and CCFs (blue) from one cell for each condition shown. Cross-correlation amplitudes are high for the positive control and wild-type CD4-Lck (top), indicating that all CD4 molecules are bound to Lck. Also shown are negative control conditions with zero cross-correlation amplitudes (bottom). Fits are shown in solid black lines. **d**. Cross-correlation quotients (*q*) for each condition normalized to the tandem control (100% co-diffusion) and Lck^CS^-Lck^CS^ (0% co-diffusion) with significant differences shown compared to the tandem (#), showing that the Lck clasp cysteines are essential for the high CD4-Lck occupancy. ‘CD4-Lck (swap)’ indicates CD4-mEGFP co-expressed with Lck-mCherry2. **e**. and **f**. Diffusion coefficients for Lck-mEGFP (green) and CD4-mCherry2 (magenta) derived from the fitted ACFs showing a significant increase in the diffusion speed of both CD4 and Lck when the Zn^2+^ clasp is disrupted. In **d, e** and **f** each circle represents a spatially averaged line-scan measurement from one cell. Cells were pooled from three independent replicates for 6-13 cells per condition. Significance testing was performed with a one-way ANOVA followed by Dunnett’s correction in **d** and Tukey’s correction in **e** and **f** for multiple comparisons. \*\*\*\**P* < 0.0001, \*\*\**P* < 0.001, \*\**P* < 0.01.

Wild-type CD4 and Lck produced a very high *q* value (96% ± 8%), irrespective of which FP was conjugated to either protein (**Figure 2d**), indicating that essentially all CD4 was co-diffusing with Lck. To rule out any effects of non-specific association, we generated three different negative controls: CD4-Lck^CS^, CD4-Fyn and Lck^CS^-Lck^CS^. In Lck^CS^, the clasp cysteines essential for binding CD4 are mutated to serine to abolish the interaction^19^. Fyn is a Src-family kinase that is structurally similar to Lck^25^ but cannot interact with coreceptors^26^. In the last condition, we expected the co-expression of Lck^CS^-mEGFP and Lck^CS^-mCherry2 to control for any lipid-mediated targeting as this form of Lck cannot homodimerize^27^ but the lipid attachment sites remain intact. No cross-correlation was detected in any of the negative controls, confirming that the high CD4-Lck occupancy was specific only to an intact Zn^2+^ clasp (**Figure 2d**).

While CD4 is a transmembrane protein, Lck is only anchored to the inner leaflet through lipid-modified residues in the SH4 domain^28^. Given the high CD4-Lck occupancy, this suggested that the diffusion of coreceptor-bound Lck would be significantly slower than free Lck according to the Saffman-Delbrück model of diffusion in biological membranes^29–31^. To test this, we used sFCCS to determine the diffusion coefficients of CD4-Lck, CD4-Lck^CS^ and CD4/Lck expressed alone (see **SI Materials and Methods** for transit time calculations). Wild-type CD4-Lck exhibited similar diffusion coefficients (Lck = 0.29 ± 0.05 µm^2^/s, CD4 = 0.21 ± 0.05 µm^2^/s), and mutation of the Lck clasp produced a significantly increased diffusion speed (Lck^CS^ = 0.97 ± 0.29 µm^2^/s, CD4 = 0.33 ± 0.07 µm^2^/s), as expected, indicating that disruption of the complex increased the lateral speed of both CD4 and Lck (**Figure 2e, 2f**). In the case of CD4, this diffusion was comparable to when the protein was expressed alone (0.39 ± 0.11 µm^2^/s). When Lck was expressed alone its diffusion speed was substantially reduced compared to Lck^CS^ (0.64 ± 0.15 µm^2^/s) and we speculate that this is due to the propensity of Lck to homodimerize through the clasp. Therefore, we concluded from these experiments that the bulk of CD4 molecules co-diffuse with Lck in the steady state, and confirmed that the Zn^2+^ clasp is necessary for this interaction to occur.

### The CD4 amphipathic helix is necessary for efficient Lck association

There is evidence from early biochemical mutagenesis studies that intracellular sequence elements in CD4 other than the Zn^2+^ clasp might contribute to the interaction with Lck^19,32^. When overlaid with the CD4-Lck NMR structure^12^, these elements appear to fall into three regions: the membrane-proximal basic-rich region (BRR), the amphipathic α-helix, and the unstructured C-terminal ‘tail’. To analyze the contributions of these features of CD4, we generated a panel of full-length CD4 molecules each bearing mutations in either the BRR, the helix or the tail (**Figure 3a**). To avoid making large truncations that might alter the spacing between the plasma membrane and the Zn^2+^ clasp, we replaced each sequence of interest with a flexible (Gly-Ser)_n_ linker of equivalent length. Each mCherry2-tagged CD4 construct was then co-expressed with wild-type Lck-mEGFP in HEK293T cells for sFCCS measurements (**Figure 3b**). We found that, while the BRR did not contribute to Lck association, mutation of the amphipathic helix reduced CD4-Lck occupancy by 27% (**Figure 3c**). When both were mutated simultaneously, occupancy was reduced by 28% indicating a significant role for the amphipathic helix in efficient CD4-Lck complex formation. As controls, we mutated the CD4 palmitoylation sites in CD4 (CD4Δpalm) which had no effect on CD4-Lck occupancy^33^ or the clasp cysteines (CD4^CS^) which abolished the interaction (**Figure 3c**). This demonstrated that an intact Zn^2+^ clasp is necessary but insufficient for the formation of high-occupancy CD4-Lck complexes. Additionally, mutation of the unstructured ‘tail’ produced a small (~14%) decrease in occupancy (**Figure 3c**). This disordered C-terminus is absent from the CD4-Lck NMR structure^12^ but may be involved in stabilizing the clasp by interacting with proximal elements of Lck. To confirm that the effects on occupancy were not due to inadvertent effects on mobility or expression, we then used FCCS to determine diffusion coefficients (**Figure 3d**) and fluorophore density (**Figure 3e**). A diminished ability to bind Lck correlated with significantly faster diffusion, as expected from our previous data, and each construct expressed at wild-type levels demonstrating no changes in expression efficiency. Overall, our mutation experiments show that the wild-type, stoichiometric interaction of CD4 with Lck relies on both the Zn^2+^ clasp and the amphipathic helix of CD4.

**Figure 3.**
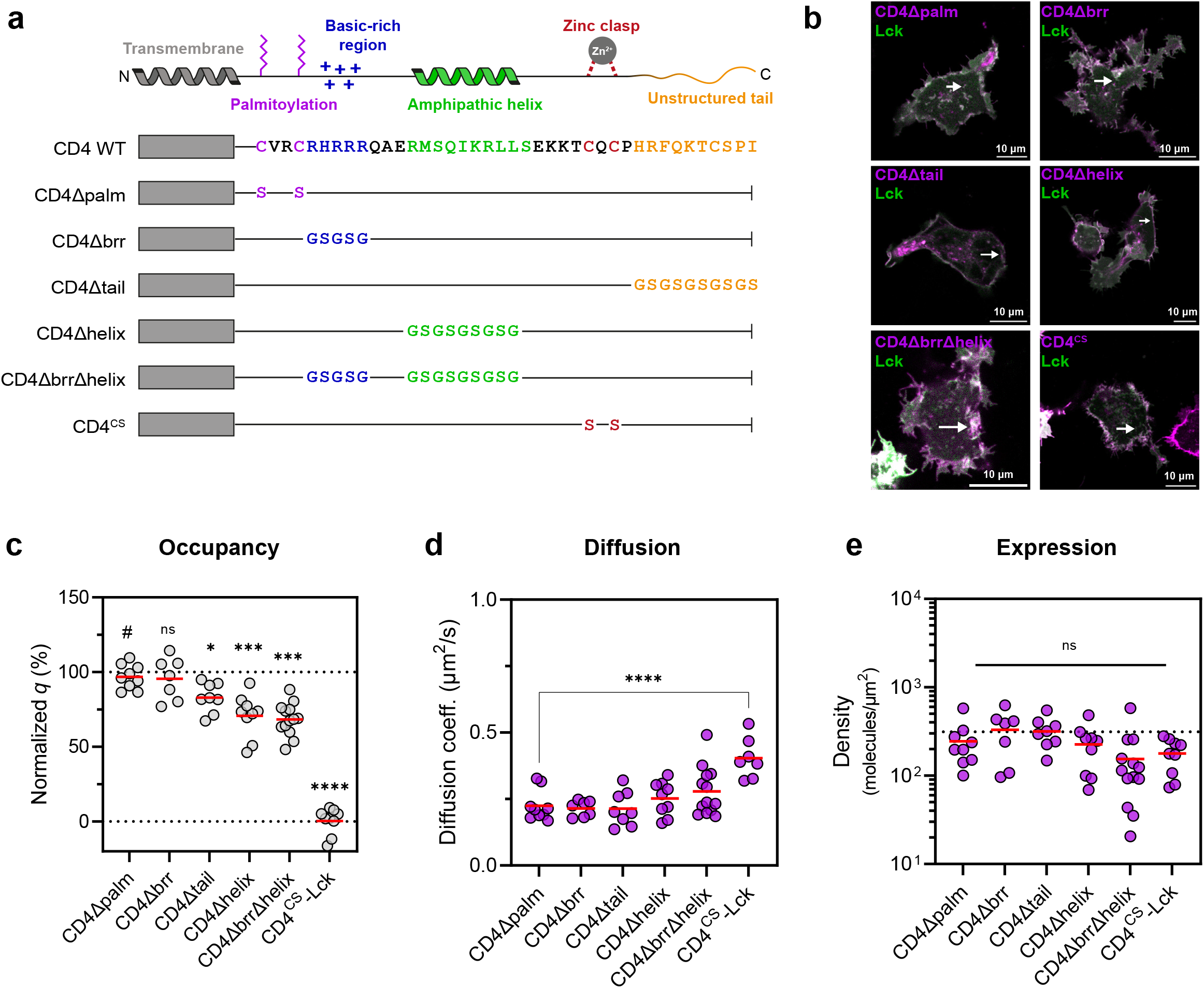
Contributions of CD4 sequence motifs to efficient Lck association. **a**. Schematic of the panel of CD4 mutants used to test the binding properties of the intracellular sequence elements indicated at the top. **b**. Confocal images of HEK293T cells co-expressing the indicated mCherry2-tagged CD4 constructs with wild-type Lck-mEGFP acquired in photon-counting mode with the white arrows indicating the approximate position of the line-scan. **c**. The CD4 amphipathic α-helix makes an important contribution to the stoichiometric co-diffusion of CD4-Lck complexes. Cross-correlation quotients (*q*) were normalized to the controls in Figure 2; statistical comparisons were made to the CD4Δpalm condition (#). **d**. CD4 diffusion coefficients are similar between constructs with an increase seen only on the loss of Lck-binding; statistical comparisons as in **c. e**. All CD4 constructs express at similar molecular densities. The dotted line indicates the average expression density of wild-type CD4 from **Figure S2b**. In **c, d** and **e** each circle represents a spatially averaged line-scan measurement from one cell, with measurements pooled from three independent replicates for 7-13 cells per condition. Significance testing was performed with a one-way ANOVA followed, in **c** and **d**, by Dunnett’s correction for multiple comparisons. \*\*\*\**P* < 0.0001, \*\*\**P* < 0.001, \*\**P* < 0.01, \**P* < 0.05.

### CD4 outcompetes CD8αβ for Lck binding in DP cells

CD8-Lck interactions are weaker than CD4-Lck interactions in co-IP assays (**Figure 1c**) and there is a two-fold difference in affinities when assayed as isolated polypeptides in solution^12^, perhaps reflecting the lack of known secondary structure in the cytoplasmic domain of CD8α. Consistent with this suggestion, we co-transfected CD8α, CD8β-mCherry2 and Lck-mEGFP into HEK293T cells and found that the CD8αβ-Lck occupancy was 59% ± 10%, similar to that of helix-deficient CD4 (**Figure 4a**). The high occupancy of both coreceptors also suggested that they might compete for Lck when co-expressed *i*.*e*., in CD4^+^ CD8^+^ DP cells. To test this possibility, we co-transfected CD4-mCherry2 with non-fluorescent CD8α/CD8β or CD8β-mCherry2 with non-fluorescent CD8α/CD4, alongside equivalent amounts of Lck-mEGFP, and measured the coreceptor-Lck *q* in each DP condition (**Figure 4b**). The average occupancies of both CD4-Lck (57% ± 20%) and CD8αβ-Lck (37% ± 20%) in DP cells were substantially reduced compared to the single-positive (SP) conditions (*cf*. 96% for CD4-Lck, 59% for CD8αβ-Lck **Figure 2d, Figure 4a**), implying that all available kinase was partitioned between the coreceptors when Lck was limiting (DP-Lck^Low^). We hypothesized, therefore, that increasing Lck levels in excess of coreceptors (DP-Lck^High^) would reverse this trend, restoring the coreceptor-Lck equilibrium to that of SP cells. These measurements were technically challenging because increased mEGFP expression led to cell brightness beyond the detection range for single-molecule fluctuations. Nonetheless, in the narrow range of cells yielding suitable fluctuation curves, increased Lck-mEGFP expression levels (**Figure S3**) yielded average occupancy values comparable to SP HEKs for both CD4 (88% ± 23%) and CD8αβ (60% ± 12%) (**Figure 4b**). This suggested that Lck expression levels could regulate coreceptor occupancy, with CD4 outcompeting CD8αβ when Lck is limiting. To test this statistically, we pooled DP-Lck^Low^ and DP-Lck^High^ cells and used regression analysis to determine the relationship between the amount of excess Lck (Lck/coreceptor density ratio) and kinase occupancy (**Figure 4c**). Although there was no statistical link between CD4-Lck *q* and the Lck/coreceptor ratio (R^2^ = 0.001, p = 0.83), CD8-Lck *q* increased significantly with excess Lck. (R^2^ = 0.38, p < 0.0001). At low Lck concentrations, CD4 outcompetes CD8αβ and it is only when the Lck levels begin exceeding both coreceptors (Lck/coreceptor ratio >= 2) that CD8αβ-Lck reaches the occupancy values observed in cells expressing CD8αβ alone. Together, our results suggest that the CD8αβ-Lck balance in DP cells might be controlled by Lck expression levels.

**Figure 4.**
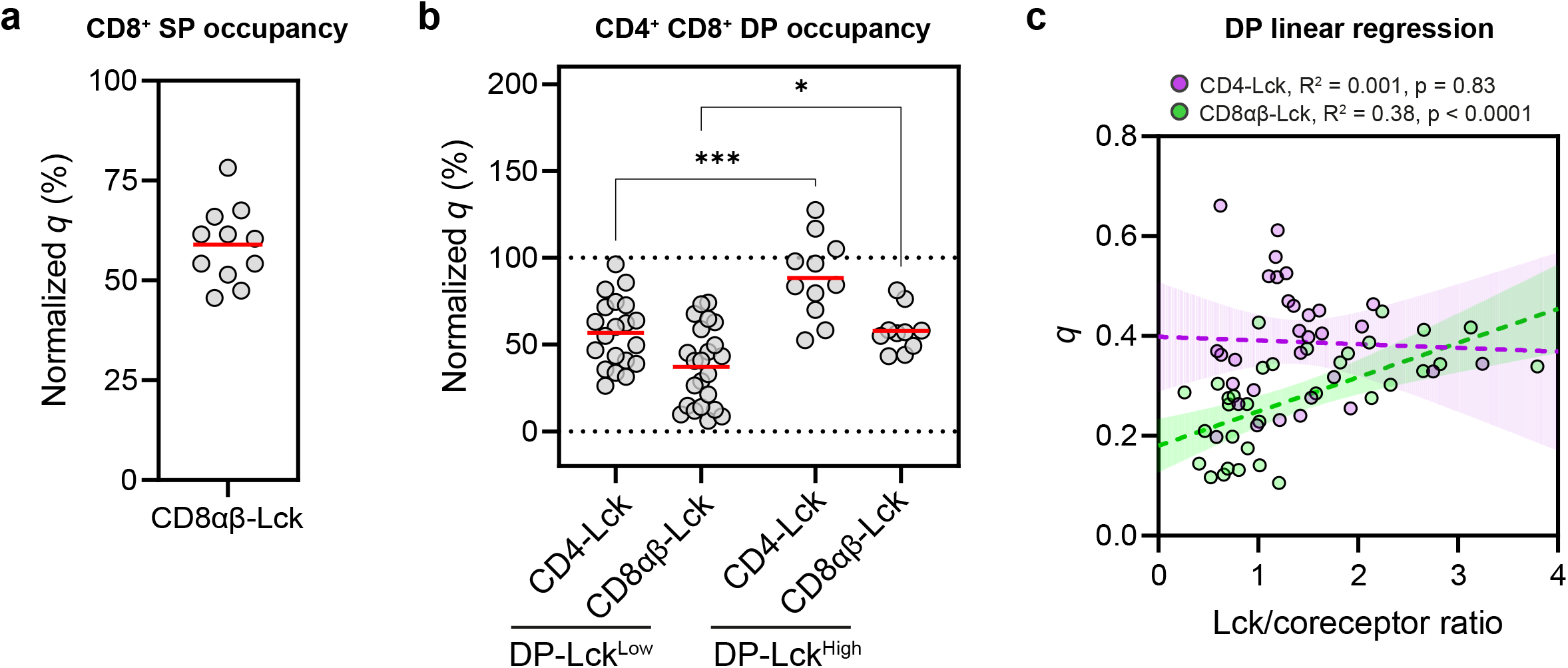
CD4 outcompetes CD8 αβ for Lck at limiting kinase levels. **a**. sFCCS measurements of CD8αβ-Lck were performed by transfecting CD8α, CD8β-mCherry2 and Lck-mEGFP into HEK293T cells and normalizing *q* values to the controls in Figure 2, yielding an occupancy of 59% ± 10% **b**. DP HEK293Ts were generated by transfecting either CD4-mCherry2 and unlabeled CD8α and CD8β, or CD8β-mCherry2 and unlabeled CD4 and CD8α. Either 25 ng or 50 ng of Lck-mEGFP plasmid DNA was co-transfected to generate the Lck^Low^ or Lck^High^ conditions, respectively. The *q* values were determined as in **a**. In DP-Lck^Low^ cells, occupancy is lower than in SP cells with a larger variance, suggesting that CD4 and CD8 compete for limiting Lck in individual cells. When Lck is in excess (DP-Lck^High^), occupancy is restored to that of SP cells indicating saturation of both coreceptors. **c**. Linear regression was used to test for a link between the amount of excess Lck (Lck/coreceptor ratio) and kinase occupancy for both CD4-Lck (magenta) and CD8αβ-Lck (green). Only the latter showed a positive, significant relationship indicating that CD4 outcompetes CD8αβ for Lck when it is limiting. F-tests were used to determine whether the slopes differed significantly from zero. Dotted lines indicate regression slopes and shaded areas indicate 95% CI. In each panel, a circle represents a spatially averaged line-scan measurement from one cell, with measurements pooled from three independent replicates for 10-23 cells per condition. Significance testing was performed with a one-way ANOVA followed by Tukey’s correction for multiple comparisons. \*\*\**P* < 0.001, \**P* < 0.05

### CD4-Lck occupancy is independent of TCR signaling

Our experiments in HEK293T cells were designed to measure *in situ* coreceptor-Lck interactions in the absence of any T cell-specific factors, but an early hypothesis for coreceptor function was that the activities of CD4 and CD8αβ could be controlled by TCR signaling^34^ potentially through regulating kinase occupancy^35^. To track coreceptor-Lck interactions in the context of TCR triggering, we transfected CD4-mCherry2 and Lck-mEGFP into Jurkat^CD4-/Lck-^ T cells (**Figure S1c**) and incubated them at 37°C on supported lipid bilayers (SLBs) functionalized with the small adhesion protein CD58. CD58 was necessary to induce Jurkat T cells to form suitably stable contact areas for sFCCS, as they would otherwise migrate or drift^36^ introducing movement artefacts in the fluctuation analysis. Then, to activate the cells through TCR engagement, anti-CD3ε (clone UCHT1) Fabs were tethered to the SLBs alongside CD58 (**Figure 5a**, ‘activated’). To control for the basal level of activation that occurs through the CD58 condition alone (‘primed’), owed to ligand-independent TCR triggering^37,38^, we generated TCR-deficient Jurkat^CD4-/Lck-^ cells (Jurkat^TCR-^) using CRISPR-Cas9 (**Figure 5a**, ‘non-stimulated’). The diminished signaling capacity of Jurkat^TCR-^ cells was verified by Ca^2+^ flux measurements when placed onto SLBs decorated with UCHT-Fabs and CD58 (**Figure S4a**). We noticed, additionally, that Jurkat^TCR-^ cells did not spread in the characteristic manner of activated T cells on SLBs (**Figure 5a, Figure S4b**). This difference was confirmed by measuring the fluorescent area (*i*.*e*., the area containing either CD4 or Lck) at the plane of contact with the SLB, and the average area was found to be significantly increased in activated condition compared to either the primed or non-stimulated setting (**Figure 5b**). CD4-Lck interactions were analyzed by performing sFCCS in the Jurkat cells that had formed stable contacts within 10 minutes of incubation (**Figure 5a**, white arrows). The cross-correlation analysis for these measurements produced high CD4-Lck *q* values, identical to the occupancies measured in HEK293T cells (**Figure 2d**), and signaling through the TCR had no effect (**Figure 5c**). This implies that CD4 is invariably bound to Lck at the T-cell surface, and that high occupancy is not an artefact of heterologous expression. These results are also consistent with our FC-IP data with Jurkat T cells expressing murine proteins (**Figure 1c**), suggesting that CD4 and Lck are coupled stoichiometrically in both humans and mice.

**Figure 5.**
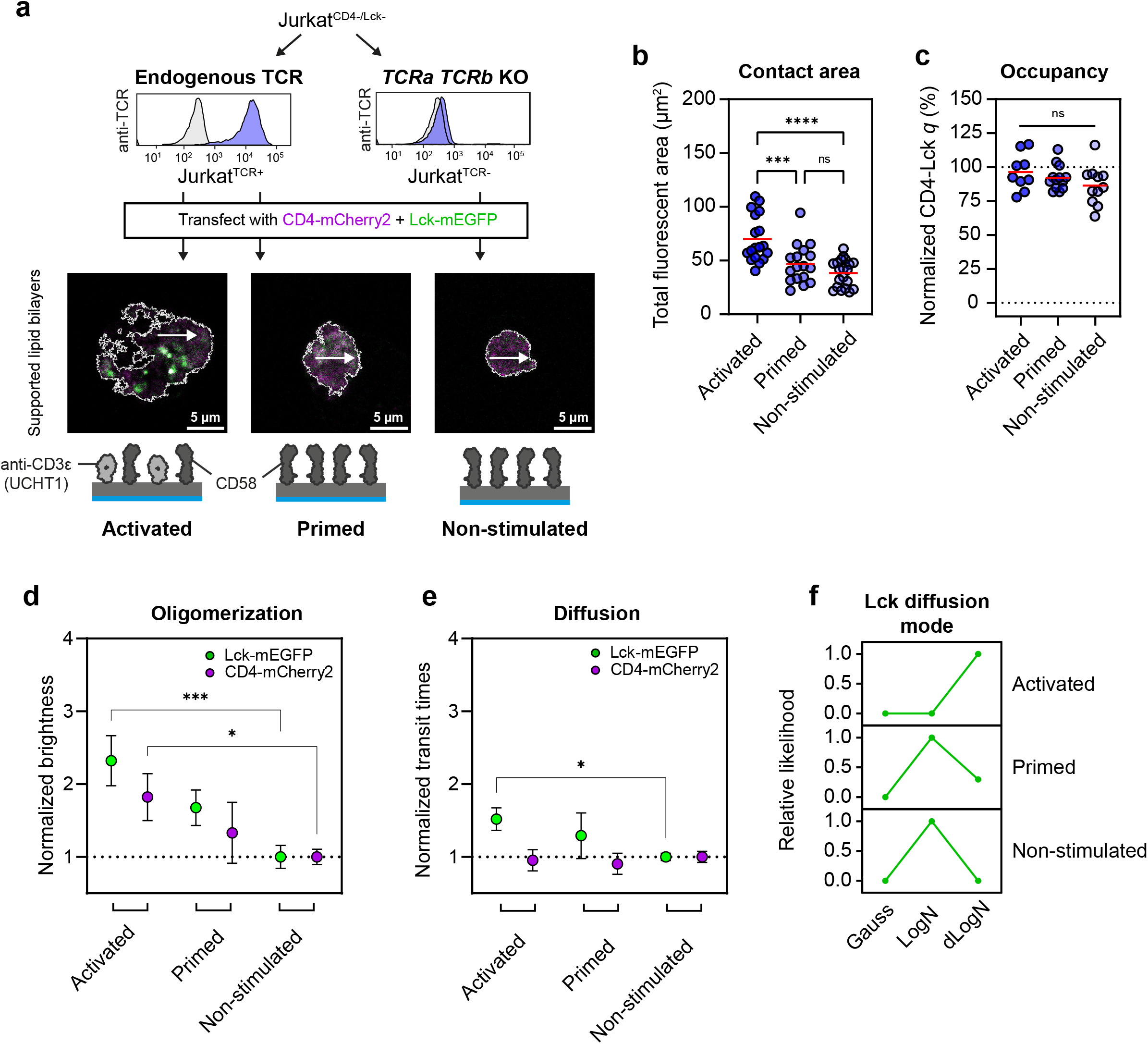
CD4-Lck occupancy is stable during early TCR signaling. **a**. TCR expression was ablated in Jurkat^CD4-/Lck-^ cells to produce Jurkat^TCR-^ cells, assayed with flow cytometry (top panel) using anti-TCR antibodies conjugated to PE (blue). Fluorescence levels of Jurkat^TCR-^ cells match the isotype control (gray), confirming knockout. Confocal images (bottom panel) of Jurkats cells co-transfected with CD4-mCherry2 (magenta) and Lck-mEGFP (green) and plated onto supported lipid bilayers (SLBs) functionalized with recombinant His-tagged human proteins. CD58 (dark gray, 200 molecules · µm^-2^) was used to anchor the Jurkat cells to the surface and anti-CD3ε Fab (light grey, clone UCHT1, 30 molecules · µm^-2^) was used for activation of the TCR. Three conditions were tested: Jurkat^TCR+^ cells on UCHT1 + CD58 SLBs (‘activated’), Jurkat^TCR+^ cells on CD58 SLBs (‘primed’) and Jurkat^TCR-^ cells on CD58 SLBs (‘non-stimulated’). The white lines indicate the outline of the cell based on fluorescence thresholding. Protein models were based on the crystal structures of a Fab fragment and hCD58 (PDB IDs: 6TCR and 1CCZ, respectively). **b**. The average spreading areas of Jurkats cells in the activated, primed and non-stimulated condition, as measured using the total fluorescent area for each cell, show an increase with TCR stimulus. Each circle indicates one cell with measurements pooled from three independent replicates for 17-20 cells per condition. **c**. CD4-Lck cross-correlation quotients (*q*) normalized to the controls in **Figure 2**, indicating that CD4-Lck occupancy does not change between conditions. Each circle indicates a spatially averaged line-scan measurement from one cell, with measurements pooled from three independent replicate for 9-11 cells per condition. **d**. and **e**. Population-level brightness (**d**) and transit time (**e**) values for the three signaling conditions, normalized to the non-stimulated setting, show an increase in oligomerization (brightness) and no major changes in diffusion speed across conditions. The median brightness/transit time was calculated for each replicate (*i*.*e*., all line-scan values for all cells on one day of measurements) and the mean value of three replicates is indicated with the circle, with error bars indicating one standard deviation. Significance testing in both **d** and **e** was performed with a one-way ANOVA followed Sidak’s correction for multiple comparisons of the same protein (*i*.*e*., only Lck or only CD4) between activation conditions. Non-significant comparisons are not shown. **f**. Statistical analysis of Lck-mEGFP transit time histograms shows that Lck diffuses freely in the primed and non-stimulated condition, but adopts a hindered diffusion mode in the activated condition only. Using a maximum likelihood estimation approach, hindrances in diffusion can be revealed by histogram fitting. The relative likelihood value for the model representing the data best is 1. Fluorescently labeled molecules diffusing freely follow a lognormal distribution (LogN) and molecules that undergo nanoscale trapping interactions result in a double-lognormal distribution (dLogN). Significance testing in **b** and **c** was performed with a one-way ANOVA followed by Tukey’s correction for multiple comparisons. \*\*\*\**P* < 0.0001, \*\*\**P* < 0.001.

T cell activation through the TCR leads to clustering of signaling proteins^39^ and single-molecule imaging experiments have shown that both CD4 and Lck can form small, nanoscale clusters within minutes of triggering^40,41^, in part through confinement in CD2 nanodomains^42^. To compare the population-level clustering of CD4 and Lck (*i*.*e*., integrated over all positions in all cells) we compared the brightness values generated from sFCS as a measure of molecular oligomerization^23^ (**Figure 5d**). Brightness appeared to increase for both CD4 and Lck upon ligation of the TCR, indicating signaling-specific oligomerization. Although no major changes were seen in the bulk transit times of CD4 and Lck (**Figure 5e**), we suspected that the diffusion mode of Lck might change. sFCS transit time histograms can be analyzed statistically using maximum likelihood estimation (MLE) to determine if the histogram is best represented by a model for free diffusion or hindered diffusion, indicating nanoscale interactions (see **SI Materials and Methods** for details)^43,44^. Using MLE analysis, we found that Lck exhibited free (lognormal/logN) diffusion in the primed and non-stimulated state and trapped (double-lognormal/dLogN) diffusion in the activated condition (**Figure 5f**). Our results indicate that early TCR signaling has no effect on coreceptor occupancy, but does lead to changes in nanoscale organization in the form of transient hindrances^42,44^ producing trapped diffusion of Lck in the activated state only.

## Discussion

The ability of CD4 and CD8αβ to recruit kinase activity to the TCR, and their importance for potentiating T cell responsiveness, first became apparent more than 30 years ago^35,45–47^. It is surprising that the precise kinase occupancy of these proteins is still controversial despite the numerical values being an important parameter of several TCR signaling models^6,48–50^. Our non-invasive sFCCS experiments showed that the coreceptors are coupled to Lck at high occupancy *in situ*, that the CD4-Lck stoichiometry relies on both the Zn^2+^ clasp and the amphipathic helix, and that kinase occupancy is unaffected by early TCR signaling. The conclusion that the coreceptors are mostly occupied by Lck was supported by our FC-IP data and is consistent with numerous early pulldown studies^8,13,14,17,51,52^. At present, we are unable to explain the recent report by Stepanek *et al*. of a much lower coreceptor-Lck stoichiometry using FC-IP^6,16^.

Although there is evidence that both CD4 and Lck can form homodimers when expressed alone^9,27,53^, our sFCCS measurements show that the CD4-Lck heterodimer dominates when they are co-expressed, consistent with earlier BRET measurements^9^. In HEK293T cells, Lck expression was reduced when it could not bind CD4, implying that the trafficking of Lck is linked to CD4 association. Lck-deficient Jurkat T cells have been shown to upregulate surface CD4 expression when transfected with an Lck transgene^54^ and pulse-chase labeling experiments indicate that CD4-Lck complexes form rapidly after translation^55^. This implies that CD4 and Lck form high-occupancy complexes prior to arriving at the cell surface. By introducing mutations into different regions of the CD4 ICD with reference to the NMR structure of the Zn^2+^ clasp, which was unavailable at the time of earlier mutagenesis experiments^12^, we could also examine the exact contributions made by the ICD to association with Lck. This confirmed the important contribution of the CD4 intracellular helix and helped to explain why CD8αβ, which lacks the helix, binds Lck more weakly than CD4. Alignment of mammal, bird, amphibian and fish CD4 homolog sequences indicates that residues in the amphipathic helix are highly conserved^7^ – second only to the Zn^2+^ clasp cysteines – suggesting considerable evolutionary pressure to preserve this functionally important element of CD4.

How do our results relate to the roles of the coreceptors when CD4 and CD8αβ are co-expressed in thymocytes prior to lineage commitment^1^? It is unclear whether the coreceptors need only to sequester all the available Lck during this stage^49,56^ (*i*.*e*., the ‘Lck availability’ model), or if it is also necessary that coreceptors bind Lck at low capacity in order to effect differential signal amplification^6,16^ (*i*.*e*., the ‘coreceptor scanning’ model). The high kinase occupancy, observed in all of our experiments, argues against the stoichiometry needing to be kept low to enhance antigen discrimination in T cells. In fact, a recent comparison of TCR-pMHC binding affinities shows that the orders-of-magnitude increase in TCR sensitivity afforded by coreceptors comes at the cost of a slight (two-fold) reduction in discriminatory power^57^. Our experiments with DP HEK293T cells demonstrate that CD4 outcompetes CD8αβ for Lck when the kinase is substoichiometric, but the impact of this on CD4 vs. CD8αβ loading depends on the amounts of Lck available. A remarkable study by the Singer group recently showed that DP thymocyte lineage fate is determined not by the nature of the coreceptor proteins, but rather by *cis* elements in the coreceptor loci that control their expression^58^. We speculate that, during the early stages of signaling before changes in *CD4* and *CD8* transcription occur, the higher CD4-Lck occupancy versus that for CD8αβ compensates for the remarkably low affinity of CD4-MHC II interactions^59^ helping it match the signals generated by CD8αβ during thymopoiesis.

To study the dynamics of CD4-Lck diffusion in the context of early TCR signaling we used CD58-functionalized SLBs to create stable, non-synaptic contacts between Jurkat T cells and the coverslip surface^37^. Although CD4-Lck occupancy was independent of triggering, protein oligomerization could be detected through increased molecular brightness, in addition to the transition in Lck diffusion mode indicating nanoscale hindrances. Stimulated emission depletion (STED)-FCS measurements have previously shown that small, membrane-anchored enzymes exhibit trapped diffusion in ‘signaling nanoclusters’^42,60^ whose formation could precede the nucleation of actin-dependent TCR microclusters in T cells^39,61^. This shows that nanoscale changes in molecular organization, which likely go unseen in conventional microscopy, cause changes in molecular organization detectable with sFCS using a confocal microscope.

Our observations help settle the question of how CD4 and CD8αβ associate with Lck in T cells, and offer additional support for the notion that the principal functions of coreceptors are to facilitate MHC restriction during thymopoiesis and to amplify receptor signaling in the periphery^4,34^. An important conclusion from our experiments with DP HEK293T cells is that differences in CD4-Lck and CD8αβ-Lck coupling are sensitive to relative Lck abundance. When Lck is limiting, CD4 has an advantage and the ratio of CD4-Lck to CD8αβ-Lck increases. CD8αβ-Lck occupancy is slightly lower in the thymocyte FC-IPs (~55%) compared to DP HEKs (~60%) while CD4-Lck is stoichiometric in both, suggesting that Lck expression levels might be regulated during thymopoiesis to tune coreceptor-dependent TCR signaling.

## Materials and Methods

See SI for a full description of all materials, methods and calculations.

## Author contributions

A.M.M., A.M.S, S.J.D., and M.L.D. designed research; A.M.M. and E.J. performed experiments; A.M.M. and F.S. analyzed data; A.M.M., S.E.F., S.J.D., and M.L.D. wrote the manuscript with input from all authors.

## Acknowledgements

We thank all members of our laboratory for providing helpful discussion and technical support, especially to Štefan Bálint, Elke Kurz, Lina Chen and Salvatore Valvo. This work was supported by the Wolfson Imaging Centre Zeiss (780i LSM) and the Kennedy Institute flow cytometry facility. FS thanks EMBO and HFSP for the long-term postdoctoral fellowship support (EMBO ALTF 849-2020 and HFSP LT000404/2021-L). AMM and SJD are funded by the Wellcome Trust (grant codes 108869/Z/15/Z and 207547/Z/17/Z, respectively). MLD is funded by the Kennedy Trust for Rheumatology Research (Cell Dynamics Platform KENN 20 21 17).

## Data availability statement

All .fcs and .lsm5 files have been deposited at https://osf.io/fumvg/?view_only=72250deaa757430a9ecc65d54c4099a6. All other data are available either in the manuscript or in the supplementary files. Analysis code is available for both the calcium analysis (https://github.com/janehumphrey/calcium) and the statistical analysis of sFCS data (https://github.com/Faldalf/sFCS_BTS).

## Notes

### Competing Interest Statement

The authors have declared no competing interest.

### Summary of Updates

Document typesetting error fixed

