## Supplementary information for "The kinase occupancy of T-cell coreceptors reconsidered"

### Materials and Methods

*Co-immunoprecipitation (co-IP)*

Carboxylate-modified latex (CML) beads (Thermo Fisher Scientific) were conjugated to anti-mCD4 (clone RM4-4, BioLegend) or anti-mCD8β (clone 53-5.8, BioLegend) according to published protocols[1]. Thymocytes were isolated from 6-week-old wild-type C57Bl/6 mice using a 70 µm cell strainer (BD Biosciences), and for each condition $10^7$ thymocytes were lysed in 50 µl of lysis buffer (1% NP-40S, 50 mM Tris pH 7.4, 150 mM NaCl, Protease Inhibitor Cocktail (Sigma-Aldrich) for 30 minutes at 4°C. For $Zn^{2+}$ chelation, cells were either pre-treated with 5 mM TPEN for 15 minutes or 5 mM EDTA was added to the lysis buffer. Lysates were then incubated with 75,000 antibody-conjugated CML beads for 3 hours at 4°C on a vertical rotating wheel. Beads were washed 5x by pelleting and resuspending in lysis buffer (200 µl) before staining with PE-conjugated antibodies against either Lck (3A5, SCBT), mCD4 (H129.19, BioLegend), mCD8α (53-6.7, BioLegend) or rat CD48 as a negative control (OX45, BioLegend) for 40 minutes at 2 µg/ml at 4°C. Residual bead fluorescence after washing was measured by flow cytometry (LSRFortessa™ X-20, BD Biosciences) and geometric mean fluorescence intensities determined using FlowJo software v10 (Treestar, USA).

*Cell lines and CRISPR-Cas9*

Jurkat E6.1 T cells and HEK293T cells were purchased from ATCC (TIB152™ and CRL-3216™, respectively). Jurkat T cells were cultured in complete R10 medium (RPMI-1640, 10% FBS, 1% penicillin-streptomycin, 2% L-glutamine, 25 mM HEPES) at 37°C, 5% $CO_2$ and were maintained at a density of 0.5-1 million cells/ml. HEK293T cells were cultured in complete D10 medium (DMEM, 10% FBS, 1% penicillin/streptomycin). All cell culture reagents were purchased from Thermo Fisher Scientific unless otherwise indicated.

CRISPR-Cas9 was used to ablate the expression of human (h) TCR, CD4 and Lck from Jurkat T cells. Guides for the hTCRα, hTCRβ and hCD4 were designed as previously described[2]. For hLck, guides were designed and selected for high specificity with minimal off target activity using Benchling (hg38 reference genome). Complementary oligonucleotides with appropriate overhangs were annealed and ligated into the LentiCRISPRv2 plasmid (Addgene #52961) using the dual BsmBI restriction sites[3]. Guides used are indicated in **Table 1**. Lentiviruses were produced by transfecting 0.5 µg of the transfer plasmid plus 0.5 µg pMDG-VSVG, and 0.5µg pCMV-dR8.91 packaging plasmids into HEK293T cells using GeneJuice® (Novagen) as described before[4]. After 72 hours, the supernatant containing viral particles was 0.45 µm-filtered and added directly to $10^6$ Jurkat cells. Three days later, cells were selected with puromycin (Sigma-Aldrich) at a concentration of 1 µg/ml in complete R10 medium for seven days and protein expression examined using flow cytometry (LSRFortessa™ X-20, BD Biosciences).

Table 1. CRISPR guides for gene ablation in Jurkat cells.

| **Target gene** | **gRNA sequence** |
|---|---|
| hTCRα (TRAV8-4*01) | GTTGCACCTCAGCAGAACCA |
| hTCRβ (TRBV12-3*01) | TATCCAGTCACCCCGCCATG |
| hCD4 | GGCAAGGCCACAATGAACCG |
| hLck | CCATTATCCCATAGTCCCAC |

mCD4 and mLck or mLck$^{CS}$ were stably expressed in Jurkat$^{hCD4/hLck-}$ cells using lentiviral gene delivery. mCD4 cDNA was obtained by PCR amplification of a splenic cDNA library provided by M. Vuong using appropriate oligonucleotide primers (see **Table 2**). mLck cDNA was a gift from J. Pettman. For mLck$^{CS}$, the C20S and C23S mutations were incorporated into a long 5' primer prior to PCR amplification (**Table 2**). All inserts were then subcloned into the pHR-SIN lentivirus transfer plasmid backbone and verified by sequencing (Source Bioscience).

Table 2. Oligonucleotide primers for mCD4/mLck cloning

| **Primer name** | **Primer sequence** |
|---|---|
| mCD4 forward | TAGTAGACGCGTGCCACCATGTGCCGAGCCATCTCTCTTAG |
| mCD4 reverse | TAGTAGCTCGAGGATGAGATTATGGCTCTTCTGCATCC |
| $mLck^{CS}$ forward | TAGTAGACGCGTGCCACCATGGGCTGTGTCTGCAGCTCAAACCCTGA AGATGACTGGATGGAGAACATTGACGTGTCTGAAAACTCCCAC |
| mLck reverse | TAGTAGCTCGAGAGGCTGGGGCTGGTACTGGC |

Human CD4/Lck constructs were transiently expressed in HEK293T, $Jurkat^{hCD4/hLck-}$ cells, or $Jurkat^{TCR-}$ cells using lipofection. First, GeneStrings (Thermo Fisher Scientific) were designed encoding hCD4 and hLck with C-terminal mCherry2 (Michael Davidson, Addgene # 54517) or mEGFP (Michael Davidson, Addgene # 54767)) tags, respectively, and cloned into pHR-SIN. GeneStrings for all of the constructs depicted in Figures 2 and 3 were then designed with flanking MluI/BamHI sites to subclone with either FP tag as required. The day before transfection, HEK293T or Jurkat T cells were plated in chambered coverslips (µ-Slide #1.5H, ibidi) at a density of $5\times10^4$ cells/well in complete media. 24 hours later, a total of 200 ng of plasmid DNA was prepared with 0.5 µl GeneJuice (Novagen) and 12.5 µl of serum-free media before addition to each well. Cells were grown for a further 24 hours before imaging to allow expression and maturation of FPs[5].

*Flow cytometry*

Flow cytometry was used to assess cells for the expression of surface or intracellular molecules. For surface labeling, $5\times10^5$ cells were washed in flow cytometry buffer (PBS, 1% FBS, 0.05% sodium azide all from Thermo Fisher Scientific) followed by staining with PE-conjugated anti-hCD4 (RPA-T4, BioLegend), anti-mCD4 (H129.19, BioLegend) or anti-αβTCR (IP26, BioLegend) at 10 µg/ml for 40 minutes at 4°C before fixation (4% formaldehyde in PBS for 10 minutes). For intracellular labeling, $5\times10^5$ cells were washed and fixed using the same procedure as for surface staining. Fixed cells were then permeabilized in intracellular staining buffer (PBS, 0.5% saponin, 1% FBS) and stained with anti-Lck-PE (3A5, SCBT) for 40 minutes at 4°C. After either surface or intracellular staining, cells were washed twice before analysis at a flow cytometer.

*Imaging and scanning-FCCS*

Fluorescence imaging and spectroscopy measurements were acquired on a Zeiss LSM 780 inverted confocal microscope (LSM780, Carl Zeiss) equipped with a 40x C-Apochromat 1.2 NA water-immersion FCS objective. Fluorescence was collected onto the hybrid GaAsP detectors (Channel S). Samples were excited with a 488 nm Argon laser (for mEGFP) and a 594 nm He-Ne laser (for mCherry2) using a 488/594 dichroic and the pinhole set to 1 AU. At the start of each day of experiments, point-FCS measurements were performed on a dye solution (20 nM Alexa Fluor 488) to calibrate the system by optimizing the counts per molecule when setting the correction collar and adjusting the position of the pinhole. Scanning (s)FCCS was performed in imaging mode by switching the ChS to photon-counting mode and data were collected at the basal cell membrane by acquiring a 52-pixel line (digital zoom 40x) at maximum scanning speed (3.94 µs dwell time and 2081 Hz line frequency) for $10^5$ cycles resulting in a total measurement time of ~48s. Files were saved as .lsm5 files and correlated externally using the open-source FoCuS_scan software[6]. The cross-correlation quotient (*q*) was calculated as the relative CCF amplitude from the fitted data[7,8], *i.e.*

$$q = \frac{G_{\mathrm{cc}}(\tau_0)}{\min\left(G_{\mathrm{mEGFP}}(\tau_0), G_{\mathrm{mCherry2}}(\tau_0)\right)}$$

Where $G_{cc}$ indicates the CCF amplitude and $G_{mEGFP}$/$G_{mCherry2}$ indicate the respective ACFs. If the fitted CCF amplitude was equal to the lower limit of the fitting parameters (0.001), the average of the first five raw CCF values was used instead.

The size of the 488 nm observation volume was calibrated using the solution of Alexa Fluor 488, which has a known diffusion coefficient of 435 µm²/s (see ref. [9]), using the equation:

$$\omega^2 = D \times 4 \times t_{xy}$$

Where ω is the beam waist radius, D is the diffusion coefficient and $t_{xy}$ is the average transit time. $\omega_{594}$ (0.266 µm) was calibrated using $\omega_{488}$ (0.235 µm) and the diffusion coefficient of the tandem mEGFP-mCherry2 construct. sFCCS measurements were performed at both 25°C (HEKs) and 37°C (Jurkats) and the temperature-dependence of D was corrected according to the equation[10]:

$$D(37℃) = D(25℃) \times \frac{T(37℃)K}{T(25°C)K} \times \frac{\eta(25℃)}{\eta(37℃)}$$

where η is the viscosity of water at the indicated temperature[11].

Fluorescent images were acquired using the same photon-counting settings as for line-scan measurements but at slower frequencies (~12s per 512 x 512 frame) to highlight cell morphologies. Images were processed with ImageJ (v1.53n, NIH)[12]. Fluorescent cell area was measured by merging the mEGFP and mCherry2 channels and thresholding with Otsu's algorithm[13] to determine the region-of-interest (ROIs). ROI areas were tabulated using the 'Analyze particles' function. Graphing and statistical analysis was performed with GraphPad Prism (v9.3).

*Statistical analysis of diffusion using MLE*

To determine the underlying nanoscale dynamics and to reveal the presence of hindered diffusion, the shape of the transit time histograms can be analyzed statistically as described before[14]. Freely diffusing molecules produce transit time distributions with a lognormal shape. The presence of nanoscale binding events cause an additional component in the transit time histograms (double lognormal shape). Using a maximum likelihood estimation approach, we calculated the most likely model for the data employing the Bayesian Information Criterion (BIC), assigning free or hindered diffusion for a given transit time histogram. The code for this analysis can be found at https://github.com/Faldalf/sFCS_BTS.

*Preparation of supported lipid bilayers (SLBs)*

Liposomes were prepared according to previous protocols[15]. Briefly, small unilamellar vesicles of DOPC supplemented with 12.5% DOGS-NTA were flowed onto clean glass coverslips affixed with adhesive six-lane chambers (sticky-Slide VI 0.4, ibidi) for 20 minutes and then washed with HEPES-buffered saline (HBS) supplemented with 0.1% human serum albumin (HSA) (Merck-Millipore) to remove excess liposomes. SLBs were then blocked with 5% BSA and 100 µM $NiSO_4$ in PBS for another 20 minutes before washing. Bilayers were then functionalized with His-tagged proteins (anti-CD3ε UCHT1-Fab at 30 molecules/$\mu m^2$ and CD58 at 200 molecules/$\mu m^2$, both produced in-house) by incubation for 20 minutes, and unbound proteins removed by a final wash with HBS/HSA. Protein densities were calculated using calibration curves of bead-supported lipid bilayers loaded with fluorescently labelled proteins, measured by flow cytometry, with reference to calibration beads with known fluorophore densities (Bangs Laboratories).

*Calcium release assay*

For calcium-triggering experiments, CultureWell 50-well silicon covers (Grace Bio-Labs) were cut and placed on piranha and plasma cleaned coverslips (25 mm, thickness no. 1.5; VWR). A 98:2 mixture of POPC-DGS-NTA-$Ni^{2+}$ vesicles was added to each well at a final concentration of 0.5 mg/ml (10 µl) and left for 1 hour at room temperature. Wells were washed by removing and adding 5 µl PBS five times before adding CD58 and UCHT1 Fab. Then, $5\times10^5$ $Jurkat^{TCR+}$ or $Jurkat^{TCR-}$ cells were resuspended in 100 µl of supplement-free RPMI with 25 µg/ml of Fluo-4 dye (Thermo Fisher Scientific) and incubated at 37°C for 10 minutes. Cells and SLBs were washed in complete R10 medium pre-warmed to 37°C immediately before use. Cells were washed twice and SLBs were washed ten times by adding and removing 5 µl complete R10 medium. Cells were gently dropped onto the SLB and imaged every second for

10 minutes using a 10x magnification objective to simultaneously acquire $10^2$-$10^3$ cells. Fluo-4 was excited using an argon 488 nm laser. Calcium imaging was performed on a Zeiss LSM880 inverted confocal scanning microscope. Cells were analyzed using a custom MATLAB code, as described elsewhere[16] (https://github.com/janehumphrey/calcium).

*Calculation of CD4 and Lck expression density from literature values*

Physiological surface densities of CD4 and Lck were approximated by converting absolute expression levels (molecules/cell) reported elsewhere. These values were obtained and averaged from studies using fluorescent beads[17–20], Western blots[21,22], or tandem mass spectrometry[23,24]. The average lymphocyte cell surface area (≈ 565 $\mu m^2$) was calculated assuming a 5 µm spherical radius and multiplied by a correction factor of 1.8 to account for surface roughness ($A = 4\pi \times r^2 \times 1.8$)[25,26]. This gave average surface densities of ~380 molecules/$\mu m^2$ for CD4 and ~630 molecules/$\mu m^2$ for Lck. The latter assumes that all Lck is surface-bound and is therefore an upper estimate.

### Supplementary Figures

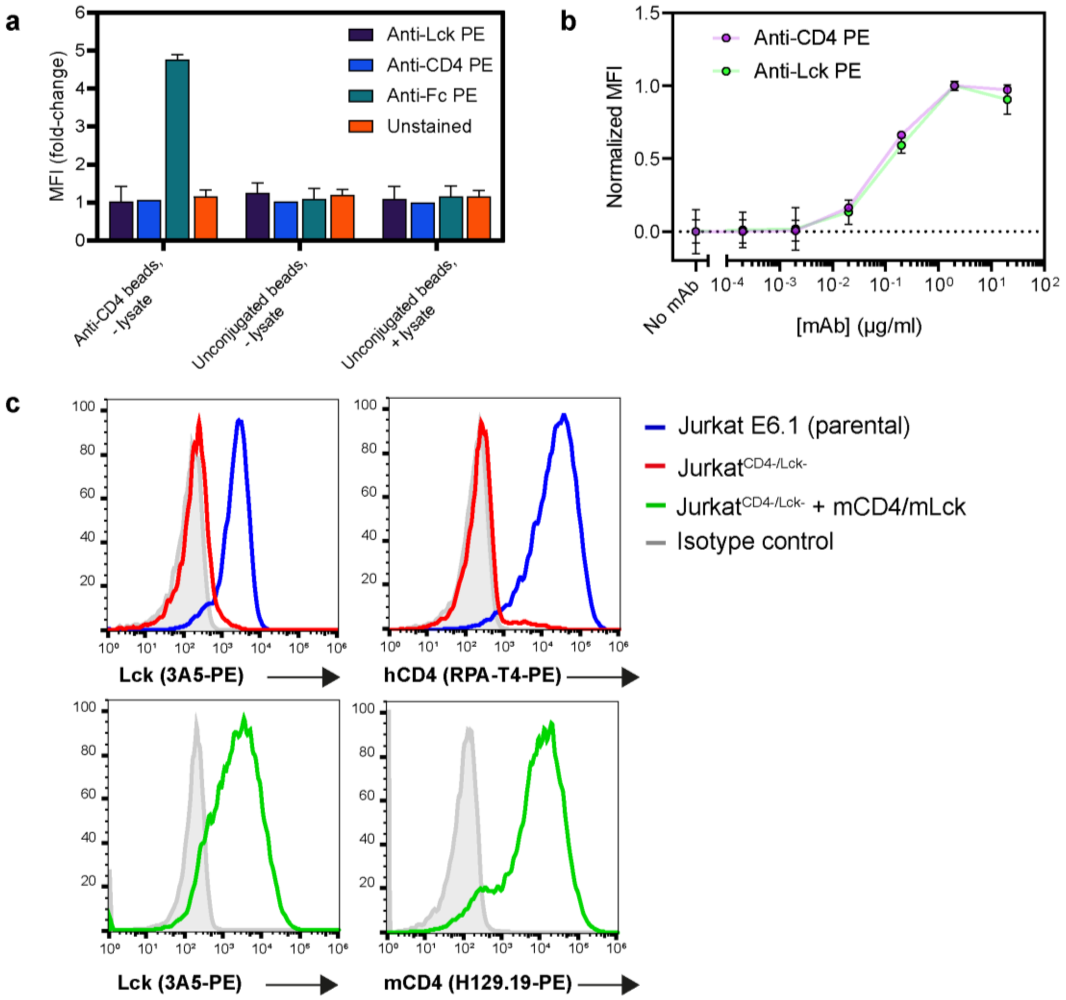


*Figure S1. Controls for the FC-IP assay.*

**a.** Controls for the FC-IP assay in which antibody-conjugated or unconjugated beads with and without thymocyte lysate incubations were stained with PE-labelled antibodies against Lck (3A5), mCD4 (H129.19) or rat IgG (MRG2b-85) and fluorescence measured compared to unstained beads. The anti-Fc labelling confirms conjugation of the capture antibody to beads. Each bar indicates the mean and standard deviation from three experiments, normalized to the lowest fluorescence intensity. **b**. Titration of the CD4 and Lck antibodies onto beads incubated with thymocyte lysates were used to determine the saturating antibody concentrations, 2 µg/ml, used for subsequent FC-IPs. Each circle indicates the mean and standard deviation from three experiments. **c.** Fluorescence histograms of the CD4/Lck-deficient Jurkat cells used for lentiviral transduction of murine CD4 and Lck.

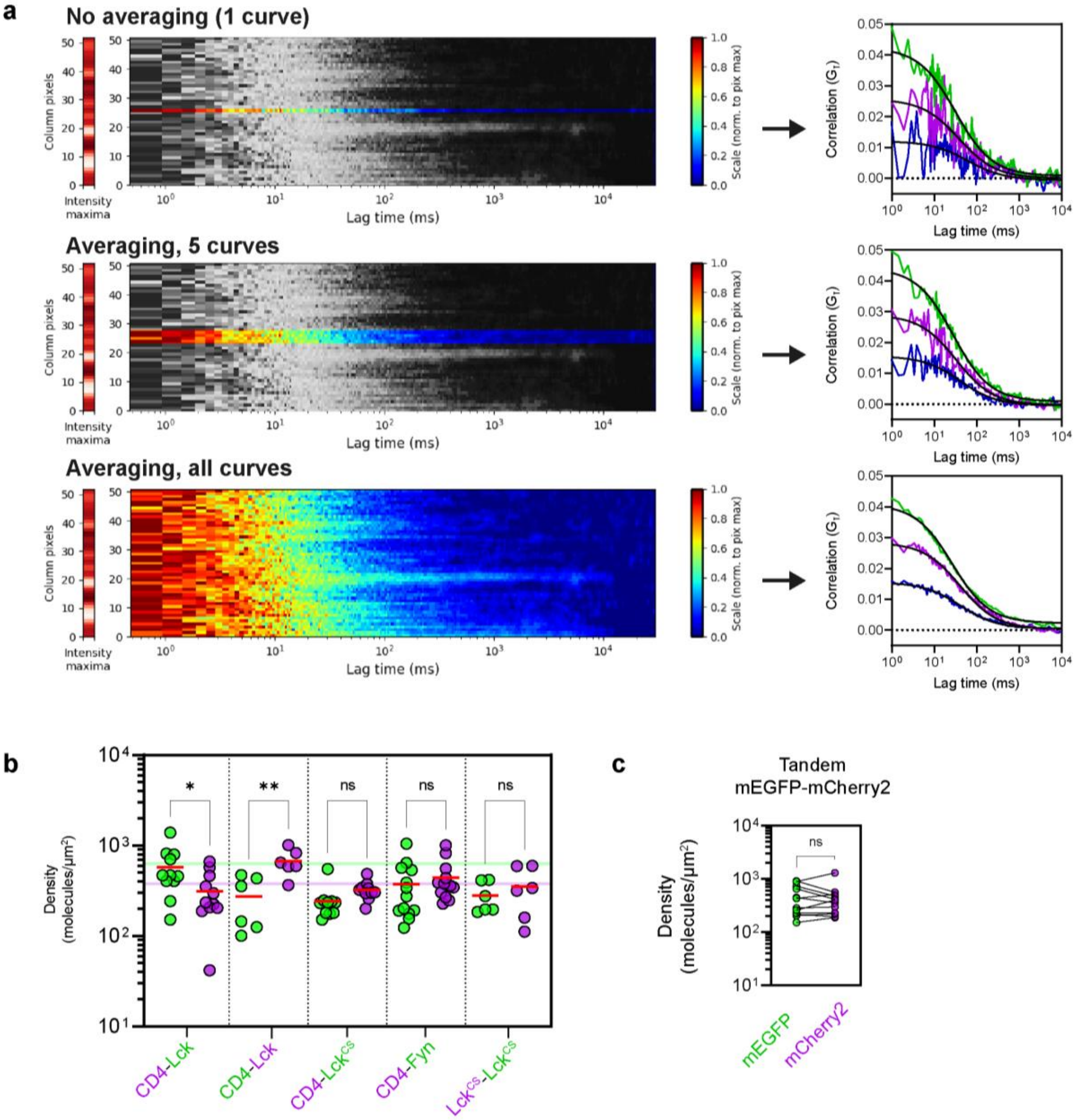


*Figure S2. Extended data for sFCCS measurements in HEK293T cells.*

**a.** Spatial averaging of tandem control sFCCS correlation carpets (left) improves the signal-to-noise ratio of the correlation curve and the quality of fitting (right). Only the carpet for mEGFP is shown while the correlation curves are shown for mEGFP (green), mCherry2 (magenta) and CCF (blue). **b.** Density measurements of the constructs measured in **Figure 2c** with green and magenta circles indicating measurements from mEGFP and mCherry2-tagged constructs, respectively. The solid lines indicate the average expression density of CD4 (magenta) and Lck (green) reported in the literature with other methods (see **Materials and Methods** for further details). Significance was assessed with a one-way ANOVA with Bonferroni's correction for multiple comparisons. **c.** Expression densities for the tandem control protein. Significance was assessed with a paired t-test.

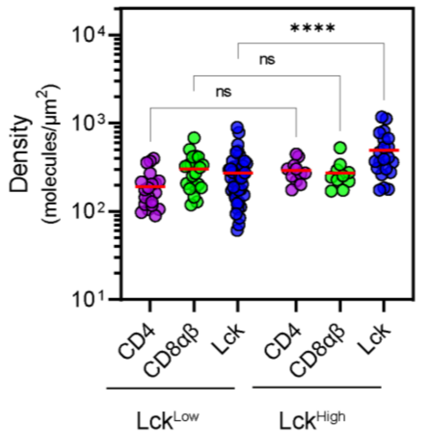


*Figure S3. Protein expression density in DP HEKs.*

Expression of coreceptors and Lck in DP HEKs with either low Lck (25 ng of transfected DNA) or high Lck (50 ng of transfected DNA). Significance testing was assessed between the same molecules expressed in either Lck$^{Low}$ or Lck$^{High}$ conditions and was performed with a one-way ANOVA followed by Tukey’s correction for multiple comparisons. ****$P$ < 0.0001.

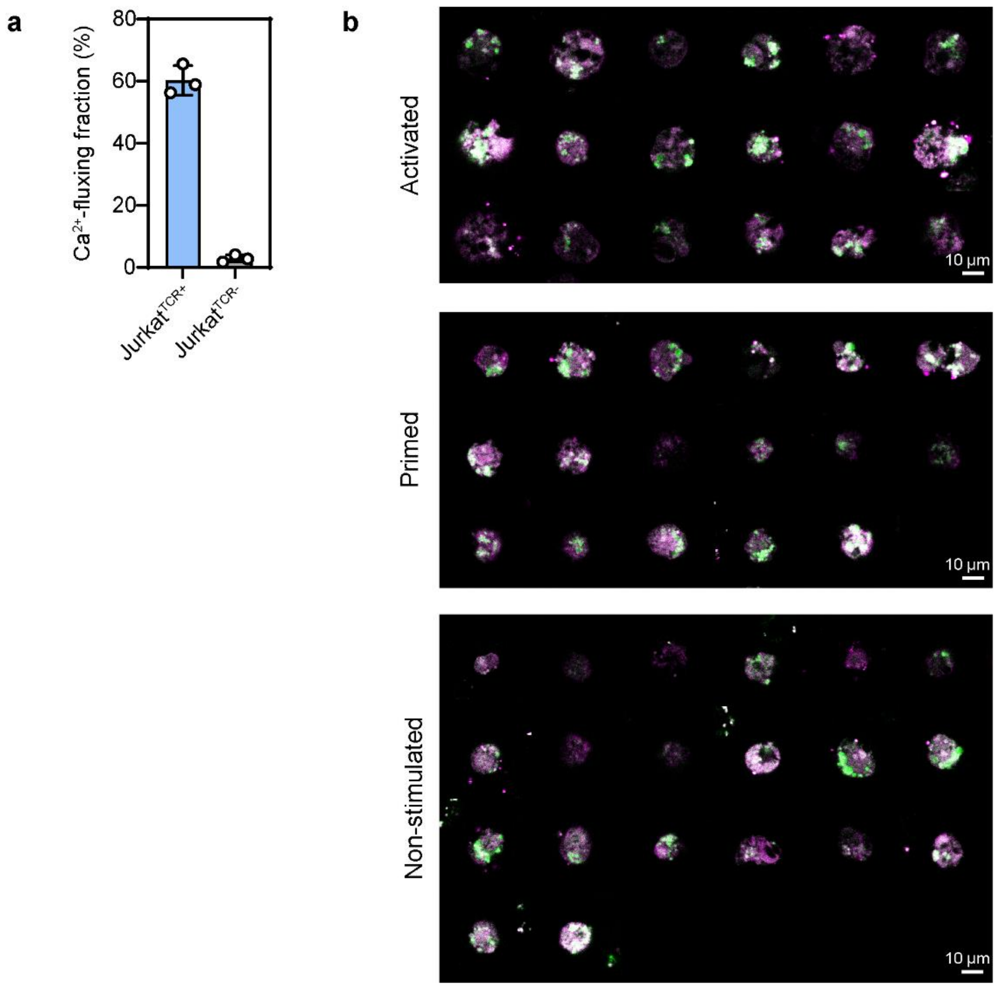


*Figure S4. Extended data for Jurkat T cell bilayer measurements.*

**a.** Jurkat$^{TCR+}$ and Jurkat$^{TCR-}$ cells were loaded with the Fluo4 Ca$^{2+}$-indicator and dropped onto SLBs presenting UCHT1 and CD58. Each circle indicates one replicate measurement of 100-1000 cells per condition. **b.** Supplementary confocal images of Jurkat T cells spreading in the different signaling conditions with CD4-mCherry2 (magenta) and Lck-mEGFP (green). Images were captured within 10 minutes of landing on the SLBs.
